# Spatiotemporal variability in transmission risk of human schistosomes and animal trematodes in a seasonally desiccating East African landscape

**DOI:** 10.1101/2023.05.25.542103

**Authors:** Naima C. Starkloff, Teckla Angelo, Moses P. Mahalila, Jenitha Charles, Safari Kinung’hi, David J Civitello

## Abstract

Different populations of hosts and parasites experience distinct seasonality in environmental factors, depending on local-scale biotic and abiotic factors. This can lead to highly heterogenous disease outcomes across host ranges. Variable seasonality characterizes urogenital schistosomiasis, a neglected tropical disease caused by parasitic trematodes (*Schistosoma haematobium*). Their intermediate hosts are aquatic *Bulinus* snails that are highly adapted to extreme rainfall seasonality, undergoing dormancy for up to seven months yearly. While *Bulinus* snails have a remarkable capacity for rebounding following dormancy, parasite survival within snails is greatly diminished. We conducted a year-round investigation of seasonal snail-schistosome dynamics in 109 ponds of variable ephemerality in Tanzania. First, we found that ponds have two synchronized peaks of schistosome infection prevalence and cercariae release, though of lower magnitude in the fully desiccating ponds than non-desiccating ponds. Second, we evaluated total yearly prevalence across a gradient of an ephemerality, finding ponds with intermediate ephemerality to have the highest infection rates. We also investigated dynamics of non-schistosome trematodes, which lacked synonymity with schistosome patterns. We found peak schistosome transmission risk at intermediate pond ephemerality, thus the impacts of anticipated increases in landscape desiccation could result in increases or decreases in transmission risk with global change.

## Introduction

Variability in environmental conditions across multiple spatial scales interacts to create highly heterogenous patterns of disease outcomes across space and time (1). For example, ambient temperature influences key life history traits of hosts and their parasites in laboratory settings (2, 3). However, in natural settings, temperature can vary substantially at hourly timescales and across microsites. Local-scale factors such as degree of habitat permanence (4–7), availability of vegetation as micro-habitat and nutrients (8), and availability of winter hardy micro-habitats (9, 10) can result in highly spatially heterogeneous host-parasite dynamics. Additionally, these dynamics are affected by seasonality, climatic cycles, and global climate change (11–13). For example, disease transmission is often elevated during warmer periods of the year (11). Disease occurrence is also typically higher in the rainy season than the dry season due to increased host activity and waterbody connectivity (14) or elevated nutrient runoff (15).

Seasonal changes in host activity and dormancy in response to cyclical climatic conditions can results in yearly peaks and troughs in transmission (11). However, different populations of hosts and parasites experience distinct seasonal fluctuations in rainfall and temperature depending on local-scale biotic and abiotic factors leading to spatially variable disease outcomes (1, 11). In addition to global changes in temperature and rainfall, human activities such as land use change can have local-scale impacts on microclimate conditions on intermediate hosts (16, 17). Considering the spatiotemporal heterogeneity in host-parasite dynamics at the local-scale, we may expect similar spatial variability in responses of hosts and parasites to global change. Anthropogenic environmental perturbations are expected to directly or indirectly increase the risk of disease incidence globally (1, 18), however, the impacts on host and parasite outcomes in regions vulnerable to drought are not extensively examined. Thus, it is imperative to better identify possible parasite outcomes in a desiccating landscape.

Schistosomiasis is a neglected tropical disease caused by parasitic trematodes in the genus *Schistosoma* and infects over 200 million people worldwide in addition to the aquatic snails that act as intermediate hosts (19). Both free-living stages of *Schistosoma haematobium,* which causes urogenital schistosomiasis, require the presence of water, yet this species occurs in African landscapes with extreme rainfall seasonality (20, 21). This seasonality results in dramatic yearly decreases in the depths and area of the ponds occupied by its intermediate snail host, *Bulinus* species, often leaving ponds dry for many months of the year. Pond ephemerality (the tendency of ponds to dry up annually) is impacted by a plethora of factors, such as pond size, depth, orientation in the landscape and human activities. The intensity of ephemerality can influence host persistence and diversity (5, 22–24). Pond desiccation forces hosts and parasites to utilize adaptive behaviors for survival (25), e.g., *Bulinus* snails undergo dormancy (known as aestivation) below the soil surface of the ponds they occupy and have an incredible capacity for population rebounding with the return of water to ponds (7, 26). However, schistosome parasite survival is greatly diminished by aestivation in laboratory studies and is understudied in the field (20). Considering the physiological challenges of aestivation on parasite survival, we hypothesize that ephemerality acts as a dampener to schistosome transmission risk.

Characterizing schistosome transmission risk with varying ephemerality can provide a blueprint of anticipated risk in an increasingly desiccating landscape. Drought intensification in East Africa is predicted with global change (27), with 10-20 million Tanzanians impacted by drought disasters and the Lake Victoria watershed of Northern Tanzania intensifying in water scarcity in the last three decades (28). To investigate the potential for pond desiccation to deter parasite transmission, we carried out a year-round evaluation of snail-parasite dynamics in ponds with varying ephemerality across six districts of the Lake Victoria watershed of Northern Tanzania. *Bulinus* snails are also frequently infected by trematodes (typically xiphidiocercaria parasites) that infect other animal species as definitive hosts, such as cattle, poultry, and wild animals. Thus, we also quantified transmission risk of these non-schistosome trematodes to assess synonymity across two parasite groups. We evaluate transmission risk in three ways: (1) snail abundance, (2) the proportion of snails infected, and (3) the total release of parasitic cercariae in the pond. Our study identifies time points and intensities of pond ephemerality that present hotspots of transmission of schistosomes and other animal trematodes.

## Methods

### Sampling sites

We surveyed 109 ponds monthly in six Tanzanian districts of the Lake Victoria watershed from August 2021-July 2022 (Figure 1). These ponds are created or modified by village communities to increase year-round water availability for the purpose of human household use (“Kisima”), for cattle use (“Lambo”), or for longer term water storage dams with unspecified use (“Bwawa”). A small number of ephemeral rivers and streams (“Mto” or “Kijito”) were also included in the study. Many of these ponds dry completely for several months of the year (Figure 2A) or dramatically decrease in size (Figure 2B) in the dry season. All ponds were chosen with approval of, and surveys were conducted in collaboration with, local village leaders. In addition, this study was conducted with permission from the Medical Research Coordination Committee of the National Institute for Medical Research (NIMR) in Mwanza (ethics approval certificate number NIMR/HQ/R.8a/Vol.IX/3462).

**Figure 1:**
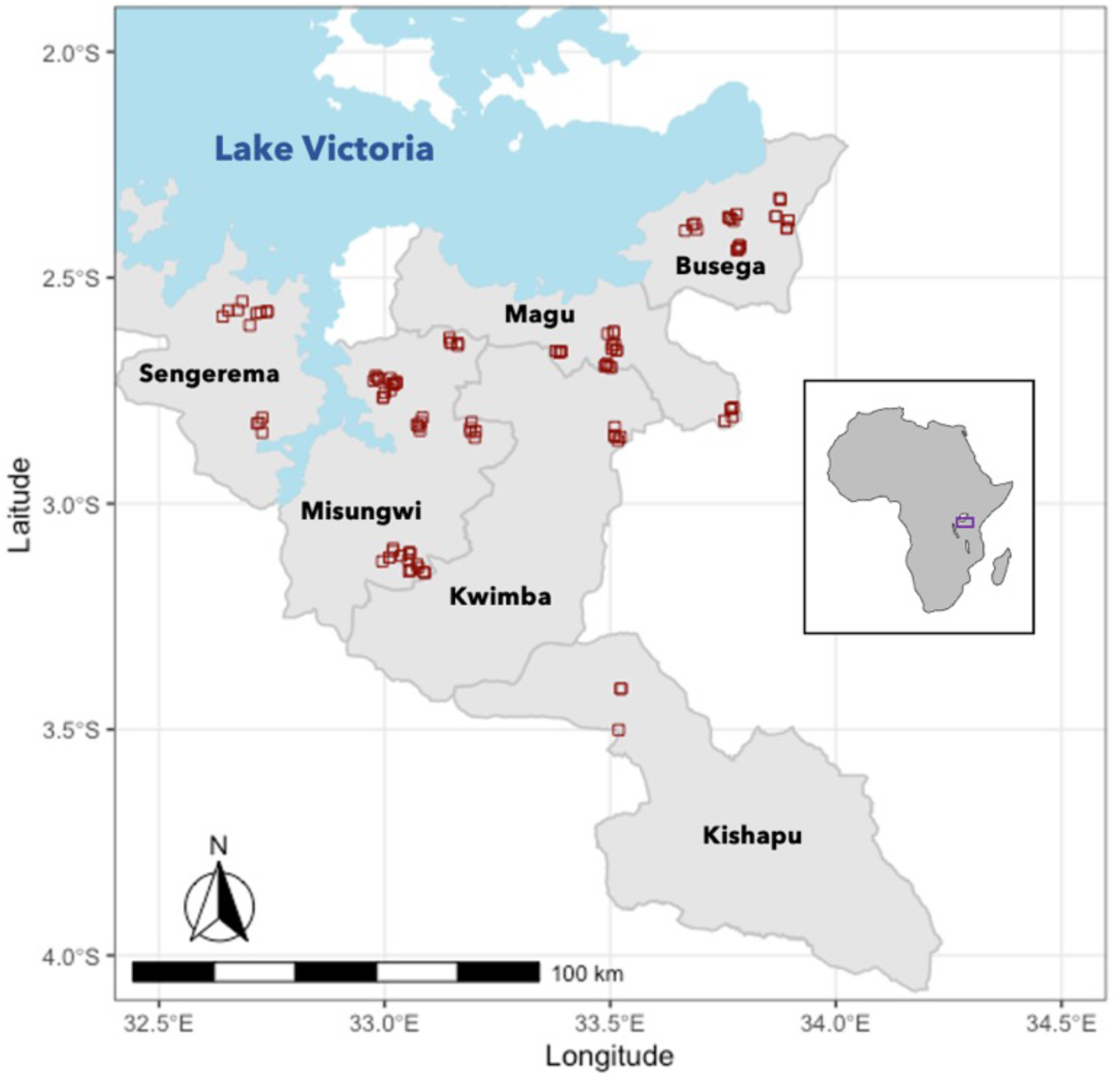
Map of localities of 109 ponds surveyed (red squares) across six districts of Northwestern Tanzania, with inset of locality of sites within the continent of Africa.

**Figure 2:**
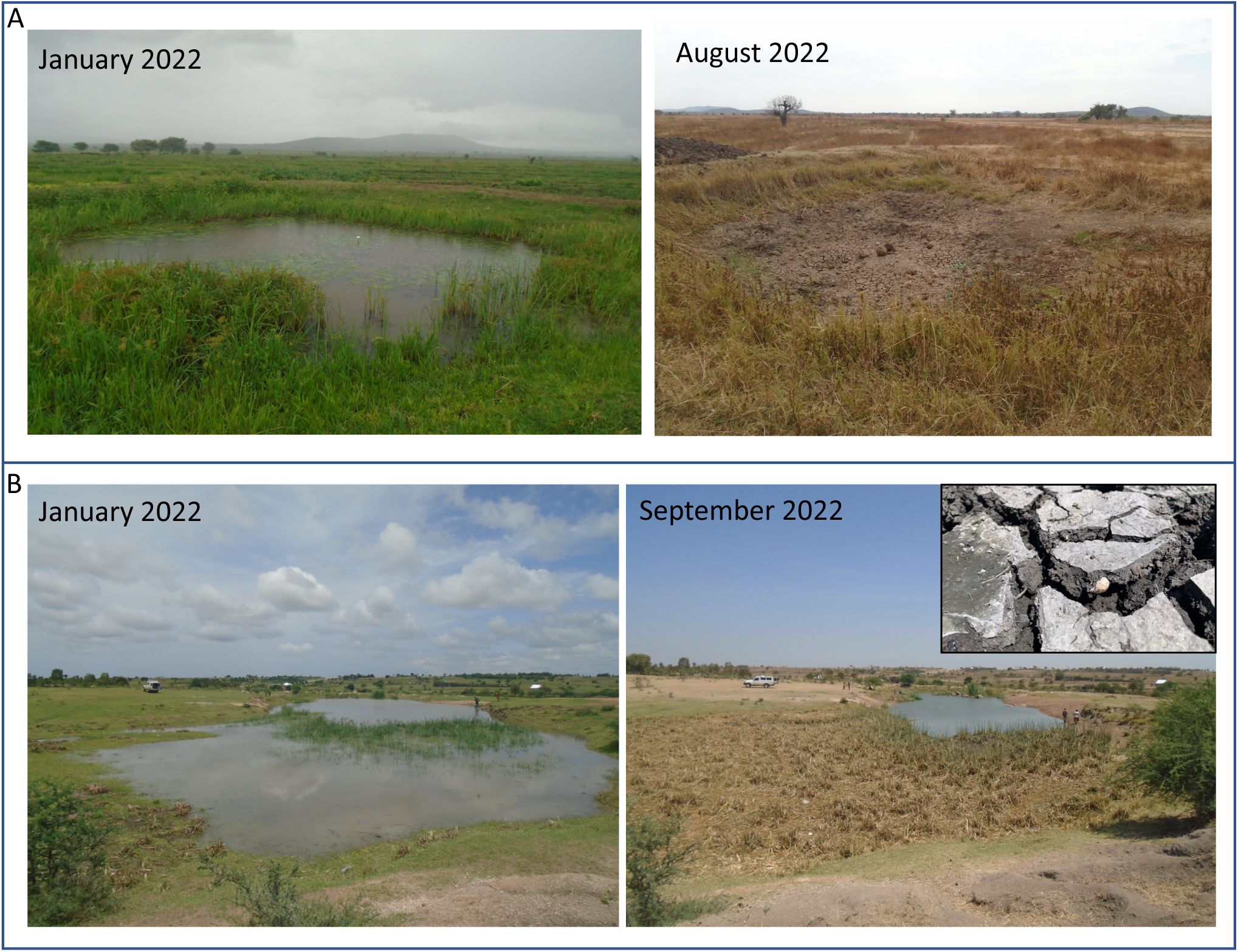
Change in area of A) a desiccating pond in Misungwi district, Kisima cha Longo (reduction of pond area of 100%) and B) a non-desiccating pond in Busega district, Lambo la Wachina (reduction of pond area of 79.78%) within a survey year. Inset shows the shallow substrate depth at which an aestivating snail was found in a pond in Busega district, Lambo la Wachina (the pictured snail had died).

The Lake Victoria watershed is typically characterized by short and long rainy seasons, *Vuli* (October-December) and *Masika* (March-May), respectively, However, a delayed *Vuli* commencing in December 2021 resulted in a combining of the two rainy periods in our sampling period (Figure 3A). Rainfall data (in mm per day) was obtained from the Mwanza weather station between 23 August 2021 and 27 July 2022 (29).

**Figure 3:**
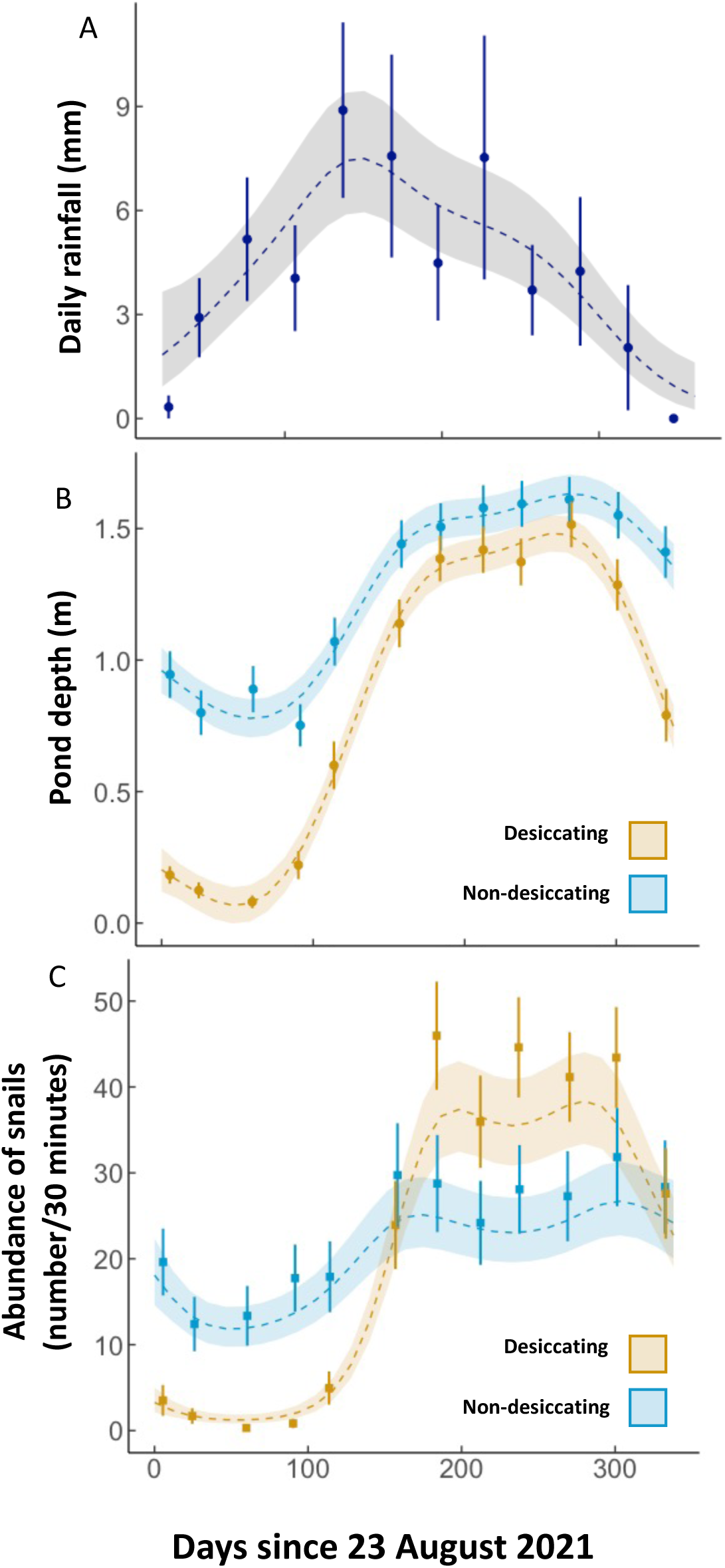
Generalized Additive Models (GAMMs) representing circannual (A) rainfall patterns (mm per day) in Mwanza, Tanzania of data from the airport weather station, and variability in B) pond depth (m) and C) *Bulinus* snail abundance (number collected in 30-minute survey) in our 109 sites. The shaded region surrounding the line represents the best fit ± 1 standard error. Points represent monthly means and standard error. Depth and snail abundance were significantly non-linear and different between the two ephemerality categories.

### Snail surveying and collection

During monthly site visits, each pond was surveyed for maximum length, width perpendicular to maximum length, and depth at center in meters. Dry ponds for which these dimensions were 0 meters were noted as such. All ponds that were too deep in the center to measure were assigned a depth of 2 meters. In addition, two researchers conducted time-constrained net sampling using metal mesh scoop nets to collect *Bulinus* snails for 15 minutes (leading to a total of 30 minutes of surveying per pond per month). Snails were placed in Nalgene containers in a cooler and brought back to the lab at the NIMR Mwanza Centre for cleaning, counting, and quantifying parasites (“shedding”).

### Identifying and quantifying parasite shedding

*Bulinus* snails were shed for patent infections in individual 30 ml beakers with 25 mL bottled water for 24 hours in natural light conditions. Following this full day shed, beakers were examined under a dissecting microscope at 10-25x for the presence of cercariae (larval forms) of schistosome and non-schistosome trematodes. The cercariae of these two groups are distinguished by size, shape, and movement (30). *Schistosoma haematobium* cannot be morphologically distinguished from *S. bovis* or their hybrids (31), therefore we represented all these individuals as “schistosomes”. Non-schistosomes were overwhelmingly represented by xiphiodiocercariae. If the presence of trematodes was confirmed, cercarial intensity was quantified after staining with Lugol’s Iodine and homogenization by gentle pipetting. For schistosomes, if the estimated number of cercariae was below 200, all cercariae were counted. If the number was larger, a subsample of 18.5% of the beaker’s bottom area was counted and multiplied by 5.412 to extrapolate for the total area of the beaker. For non-schistosome trematodes, only the subsample approach was taken due to higher intensities being typical.

### Identifying potential infected aestivators

We identified if snails carried infections through aestivation using the timing of infections relative to emergence from aestivation. The prepatent period that is necessary for infections to develop until cercariae release typically take 6-18 weeks in a laboratory setting for *Schistosoma haematobium* (32). In addition, *Bulinus* species typically aestivate in the upper periphery of ponds (21), necessitating ponds to fill for aestivating snails to emerge from the soil and be receptive to miracidia once the water has returned to ponds. This could take just a few days or several weeks from the onset of rain, depending on the size of the pond. As a result, we infer that infections that were detected less than 60 days following the last dry survey were acquired before aestivation, with increased confidence in those detected <30 days after the last dry survey.

### Statistical analyses

We utilized Generalized Additive Mixed Models (GAMMs) in the R package mgcv to evaluate how water depth (Gaussian distribution), snail abundance (Quasipoisson distribution), and infection prevalence (binomial error distributions) and total cercariae per pond (Quasipoisson distributions) of the two parasite groups varied over the course of a year. GAMMs are effective at evaluating smoothed, non-linear relationships over time. Thus, we represented the annual trend as a smooth term. For Gaussian and binomial error distributions, we fit models with Restricted Maximum Likelihood (REML), whereas as for Quasipoisson models, we fit with Quasi-Penalized Likelihood (33). We fit models with continuous autoregressive-1 error structures to account for repeated measures, except for in prevalence models due to failed convergence. For all models, we fit the dynamics of non-desiccating waterbodies with a reference temporal smooth and tested for significantly different dynamics in desiccating waterbodies with a temporal difference smooth (33). Lastly, we included pond as a random effect in the GAMMs to account for nonindependence in the monthly repeated observations from these replicated sites. We also ran Generalized Linear Models (GLMs) with the R package glmmTMB with binomial error distributions on all snails collected from each pond to assess if cumulative yearly infection prevalence varied among the two ephemerality categories.

As ephemerality can also be characterized beyond a simple dichotomy of ponds that fully desiccate or not, we also ran similar binomial GLMs to assess if yearly transmission risk differs with intensity of ephemerality (% pond reduction in a year). We summed the total number of infected and uninfected snails per pond per year for each parasite group to evaluate cumulative yearly parasite transmission risk. We then calculated the surface area of each pond on each visit by assuming an elliptical shape (Surface area = 1/2length*1/2width*π) and the percentage size reduction of each pond ((max-min)/max*100) as a measure of ephemerality intensity.

## Results

We collected a total of 30,137 *Bulinus* snails across 12 monthly surveys of 109 ponds, of which 482 snails were infected with schistosome trematodes (1.6%) and 1,592 snails were infected with non-schistosome trematodes (5.28%). Ponds were distinguished as desiccating (n=59) or non-desiccating (n=50) depending on whether water was completely absent for at least one monthly survey (Figure 2). While non-desiccating ponds had water year-round, water levels contracted between 39.46 to 99.97% the area of these ponds at their lowest observation (Figure 2B), potentially forcing snails to aestivate due to changes in water temperature, depth and quality (20). Non-desiccating ponds varied in depth through the survey period (reference smooth, p<0.001). The two pond types varied in their seasonal patterns in depth (difference smooth, p<0.001), with a more dramatic seasonal variability in desiccating ponds over non-desiccating ponds (Figure 3B).

Host and parasite dynamics were highly variable across the circannual cycle (August 2021-July 2022), with differing patterns between desiccating and non-desiccating ponds. *Bulinus* abundance number closely mirrored water depth seasonality (Figure 3C), with non-linear patterns of abundance over time in non-desiccating ponds (reference smooth, p=0.003) and a more dramatic boom and bust pattern in desiccating ponds over non-desiccating ponds (differential smooth, p<0.001). While desiccating waterbodies had a potentially elevated transmission risk due to rapidly growing snail population numbers following the onset of rain, non-desiccating waterbodies had a subdued but more constant snail abundance.

Schistosome infection patterns (Figure 4A) indicate a non-linear ebb and flow in non-desiccating ponds in terms of prevalence (reference smooth, p<0.001), and consistent, significantly lower peaks in non-desiccating ponds after about day 50 (differential smooth, p=0.044). A similar non-linear pattern is seen of schistosome cercariae release in non-desiccating ponds (reference smooth, p<0.001) with significantly lower peaks in desiccating ponds (differential smooth, p=0.014). In both pond types, there are two primary peaks following the onset of rains (peak rainfall ∼150 days); one in mid rainy season and another early in the dry season. Yearly risk of transmission of human schistosomes is substantially lower in desiccating ponds than non-desiccating ponds, with snails being 4.6 times more likely to be infected in the latter (binomial GLM; p<0.001).

**Figure 4:**
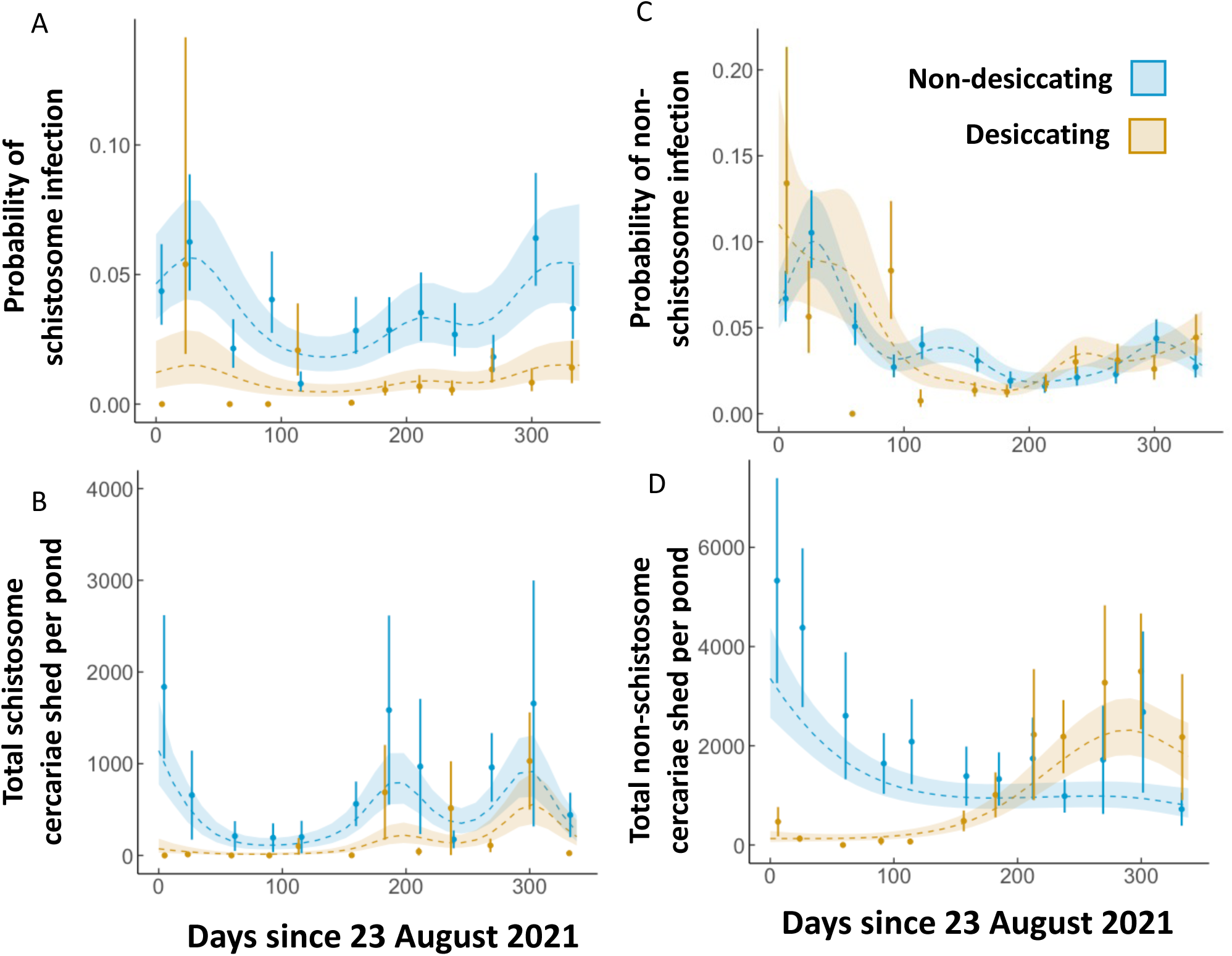
GAMMs representing circannual A) schistosome prevalence, B) schistosome intensity, C) non-schistosome prevalence, and F) non-schistosome intensity in desiccating and non-desiccating ponds. The shaded region surrounding the line represents the best fit ± 1 standard error. Points represent monthly means and standard error. All patterns were significantly non-linear and different between the two ephemerality categories.

The non-schistosome trematode GAMMs indicate significantly non-linear seasonal patterns of infection prevalence and cercarial release in non-desiccating ponds (reference smooths, p<0.001, Figure 4C-D). Non-desiccating ponds had an early-mid rainy season peak of infection prevalence whereas the rainy season infection peak of desiccating ponds is 3-4 months later (differential smooth, p<0.001, Figure 4C). In addition, there is a large peak of cercarial release only in desiccating ponds, just preceding the next dry season (differential smooth, p<0.001, Figure 4D). *Bulinus* snails are 1.6 times more likely to be infected by non-schistosome trematodes in non-desiccating ponds than desiccating ponds (binomial GLM; p<0.001), which is a substantially smaller difference than schistosomes.

Cumulative yearly prevalence varied considerably with intensity of pond ephemerality (% pond reduction in the dry season) for both parasite groups. Yearly schistosome infection had a significantly non-linear relationship with ephemerality, where prevalence peaked in ponds at intermediate ephemerality (maximum at ∼80% reduction in area, Binomial GLM; p<0.001, Figure 5A). Yearly non-schistosome infection prevalence peaked in ponds with lower ephemerality (∼50% reduction in area) and decreased steadily with increasing ephemerality (Binomial GLM; p<0.001, Figure 5B).

**Figure 5:**
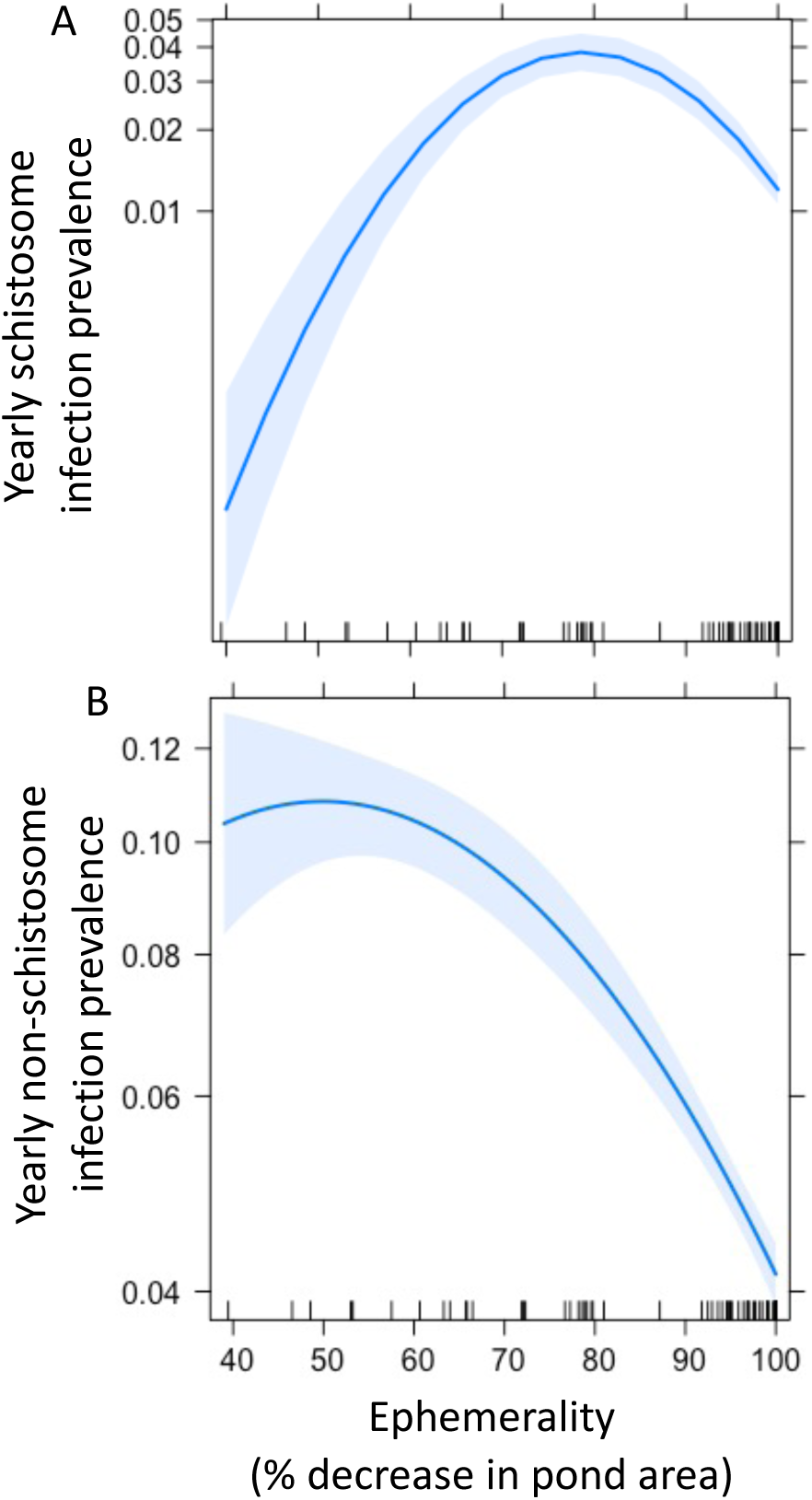
Overall yearly A) schistosome and B) non-schistosome trematode infection prevalence in across an ephemerality gradient (% decrease in pond area in the dry season). Schistosome prevalence peaks at intermediate ephemerality (∼80% area decrease, Binomial GLM, p<0.01). Non-schistosome prevalence peaks at low ephemerality (∼50% area decrease, Binomial GLM, p<0.01).

A small number of snails infected with either schistosomes (n=5) or non-schistosome (n=15) trematodes were identified less than 60 days following a survey where the pond was dry (Table 1). This provides evidence that snails do aestivate while infected, albeit rarely, and emerge and shed parasites.

**Table 1:** Case studies of infections identified <60 days after ponds were dry. Information is provided on parasite type, pond location, number of concurrent dry surveys of ponds prior to infections, number of snails in survey with first infections, number of infected snails in survey with first infections, and number of days before infections were detected since pond was last identified as dry.

| Parasite type | Pond name, Village, District | Number of dry surveys | Number of snails in survey | Number of infected snails in survey | Number of days since last dry survey |
| --- | --- | --- | --- | --- | --- |
| Schistosome | Lambo la Mhana, Ngudama, Misungwi | 1 | 100 | 1 | 31 (1 survey prior) |
|  | Lambo la Nzego Matunge, Shilalo, Misungwi | 3 | 14 | 1 | 54 (2 surveys prior) |
|  | Lambo la Joseph Malyengete, Isole, Sengerema | 3 | 10 | 3 | 54 (2 surveys prior) |
| Non-schistosome | Kwa Fumu, Nyang'holongo, Misungwi | 1 | 3 | 1 | 21 (1 survey prior) |
|  | Lambo la Sospeter Walwa, Ikungumhulu, Misungwi | 1 | 32 | 13 | 32 (1 survey prior) |
|  | Lambo la Nzego Matunge, Shilalo, Misungwi | 3 | 3 | 1 | 32 (1 survey prior) |

## Discussion

All sites in a seasonally desiccating landscape are not equal and this leads to a spectrum of dormancy conditions for their occupying populations. Global change is impacting the timing and intensity of these dormancy periods with crucial impacts on populations during active periods of the year’s cycle. For example, Penczykowski et al. (10) demonstrated that overwintering dormancy conditions have become less harsh with increasing winter temperatures, resulting in higher plant-fungi prevalence in the springs that follow. East Africa is expected to experience increased desertification with climate change (27), with variable possible outcomes on human schistosome geographic distribution (34) and transmission potential (35) at large spatial scales. At a more local scale, Mutuku et al. (36) demonstrated that a decade-long drought resulted in a near elimination of *S. haematobium* in a Coastal Kenyan village, suggesting that the transition from moderate to extreme ephemerality could interrupt transmission cycles. With increasing drought risk, ponds are likely to experience longer dormancy periods and shorter hydroperiods (length of time a water body has standing water). Thus, with anticipated increased drought frequency and intensity, we might expect to see an interruption of the transmission cycle of human and other waterborne parasites.

In our study, peak schistosome transmission risk was seen at intermediate intensities of ephemerality which could result in highly variable future outcomes depending on the pond desiccation patterns. As ponds increase in ephemerality with drying climates, they could have higher or lower transmission risk depending on their natural tendency for drying. In the case of non-schistosome parasites, increasing ephemerality is correlated with decreasing cumulative infection risk (Figure 5B), suggesting that transmission may not be well sustained in an increasingly desiccating landscape. However, drying ponds experienced a peak in non-schistosome cercarial release in the early dry season before desiccating fully (Figure 4D), which could counter drops in prevalence if cattle and other animals continued to use water at low depths. This large non-schistosome cercarial peak could be evidence of reproductive compensation on the part of the parasite with the onset of seasonally harsh environmental conditions as water levels drop (37). In addition, schistosome transmission risk peaks twice following the onset of rain synchronously across both pond types, regardless of ephemerality (Figure 4A-B), the first likely associated with infections persisting through aestivation and second likely a result of new infections following the onset of rain. Seasonal transmission patterns of non-schistosome parasites, on the other hand, are highly asynchronous across the two ephemerality categories (Figure 4C-D). This highlights the importance of long-term studies for the repeatability of infection peaks and troughs, and an understanding of their underlying predictors, especially with global change.

As seen in previous studies (7, 26), pond ephemerality was not a deterrent to *Bulinus* snail populations which have an impressive capacity for population rebounding in desiccating ponds (Figure 3c). This could be the result of factors such as escalated feeding behavior and reproduction of snails emerging from aestivation (38) and their populations being regulated by a more diverse community of competitors and predators in non-desiccating ponds than could be supported in the harsh habitat seasonality of desiccating ponds (22). While the potential for recovery of intermediate host snails following aestivation is clear, their aestivation ecology and impact on infection is still largely understudied, especially in the field (20). Aestivation imposes physiological constraints on snails (39) and infection enhances physiological stress (40), likely explaining lower infection rates in desiccating than non-desiccating waterbodies. Snails infected with schistosomes or non-schistosome trematodes can emerge from aestivation alive, though this was rare and only documented in ponds that were dry for 1-3 months (Table 1). The timeframe of our circannual cycle (beginning in the dry season) prohibited our capacity to accurately evaluate if the total duration of pond desiccation impacted host and parasite outcomes, which remains a line of inquiry for the future.

Another possible determinant of the disparity among ponds is definitive host use. Shorter hydroperiods limit human exposure to and contamination of the water, disrupting the transmission cycle. Even if snails were to get infected, they have a limited amount of time develop patent infections and thereafter, survival is limited if snails undergo aestivation with infections (40). It is, thus, surprising that ponds with the longest hydroperiods have low human schistosome transmission risk (Figure 5C). Ponds with the longest hydroperiods are those with the biggest area, such as dams. Humans likely use these less often for high transmission risk activities, such as children playing and swimming. However, these larger ponds are often used by cattle in large densities, for activities such as cattle washing stations, resulting in peak risk for non-schistosome trematodes. Non-schistosome parasites are in general at higher infection prevalence and intensities across space and time (Figure 4 & 5). Cattle interact with ponds at higher densities and are far more likely to urinate/defecate in and around ponds than humans, any time of year and regardless of the water depth. In addition, while humans in this region are enrolled in regular drug programs, often with yearly doses of the anthelmintic praziquantel, local veterinarians stated that cattle in the Lake Victoria region are provided anthelmintics on a case-by-case basis. Anthelmintics are often variably effective with increasing parasitic infection and timing of infection and we might expect cattle to have higher exposure and intensity of infections due to their indiscriminate water use and contamination, and densities.

This still leaves an open question as to why ponds with intermediate ephemerality favor schistosome transmission. These ponds do have longer hydroperiods than desiccating ponds, which perhaps creates periods with concentrated exposure/contamination risk as water depths lower in the dry season. Alternatively, snails have been observed to aestivate in non-desiccating ponds as standing water level decreases, likely triggered by unfavorable water conditions (20). However, the presence of standing water may create gentler aestivation conditions in the soil, limiting mortality of infected snails when compared to desiccating ponds. Experimental approaches and field observations may help elucidate the mechanisms underlying elevated parasite success in these intermediate pond types.

Increased ephemerality of intermediate host habitat with global change may dampen the transmission risk of waterborne parasites resulting in potentially beneficial outcomes for human, livestock, and wildlife diseases. However, it is hard to predict how all parties of these disease transmission cycles will respond to the desiccation of their landscape. Short generation times in trematodes and snails may result in adaptation to longer dormancy and shorter active periods, such as hardier dormancy phenotypes and faster reproductive cycles. Species may also experience shifts in geographic ranges in response to a changing climate, for example another human schistosome species (*S. mansoni)* has been detected at higher elevations than previously recorded in Uganda (41).

Humans have shown a history of largely small-scale mitigation to a desiccating landscape, with variable responses by different stakeholders and at different scales (28). The creation and periodic modification of these ponds, for example, was for the purpose of improving year round waterbody availability due to a history of droughts (42), as well as to provide water for an increasingly irrigated agricultural sector in Sub-Saharan Africa (43). Thus, further droughts could result in the creation of more such ponds or enlarging of existing ones to increase year-round water supplies. Either outcome has the potential to provide habitat that favors schistosome and animal trematode infection risk. Alternatively, with increasing human populations there may not be sufficient space, and this may provide an opportunity for alternative water storage and conservation methods, which may also be beneficial for human public and environmental health. Some methods used include terracing, rainwater harvest tanks, sub-surface storage and afforestation (28). In the meantime, lessons from this natural laboratory of varying ephemerality could be used to mitigate infection risk. Ponds drying out completely reduces the transmission risk of schistosome infection (Figure 4a). Seasonal pumping of water from these types of ponds to above ground water sources to dry out the soil may intensify aestivation conditions like in desiccating ponds and limit parasite survival. Additional disturbance of soil conditions, such as sun-drying, ploughing, and tilling, may further limit survival capacity (25, 44). Any such interventions should only be made with conscientious dialog with village members as to not disrupt their water practices and availability in the dry season.

Highly ephemeral waterbodies have the potential to disrupt the transmission cycle of human and other animal trematodes. Factors such as pond size, depth, shape, substrate, nutrient availability, vegetation density, and seasonal use by definitive hosts interact with a rapidly changing climate to determine infection outcomes. The dissimilar pattern of transmission risk between these two parasite groups across space and time asserts the necessity of taking a One Health approach to identifying specific mechanisms underlying their infection success in a desiccating landscape.

## Acknowledgements

We thank the field team for the collection of data that made this manuscript possible; Rashid N. Juma, Revocatus R. Silayo, Bahati Lukomeso, Amiru Swaibu, Revocatus Alphonce, James Kubeja, Kagaruki Francis, and Dennis Byakuzana. We acknowledge the support from the regional authorities of Mwanza, Simiyu, and Shinyana, as well as the leadership of communities and villages in northwestern Tanzania where surveys were conducted.

## Funding

N.C.S. and D.J.C. were supported by the US National Institute of Allergy and Infectious Diseases R01 AI50774-01. All fieldwork expenses were also funded by this source.

## Conflict of Interest

The authors have no conflict of interest to declare.

## Notes

https://doi.org/10.5061/dryad.mpg4f4r4t

### Competing Interest Statement

The authors have declared no competing interest.

